# Acute obstructive uropathy in male C57Bl/6 mice after general anesthesia with fentanyl, midazolam and medetomidine

**DOI:** 10.1101/2025.02.12.636996

**Authors:** Damian J. Wallace, Carl Holmgren, U. Czubayko, Jeanine Klesing, Daniela Martin Machado, Jason N. D. Kerr

## Abstract

The selective alpha-2 adrenoreceptor agonist medetomidine is commonly used as a component of combination solutions for general anesthesia in rodents and small animals. However, it has also been reported to increase the risk of obstructive uropathy in male mice. Here we report that in a recent cohort of mice undergoing two sequential surgical procedures using the anesthetic combination of fentanyl, midazolam and medetomidine (FMM) for general anesthesia, 60% of the male mice developed severe obstructive uropathy within 48 hours of the second surgery. We also report one case of an older male mouse developing severe obstructive uropathy after a single general anesthesia with FMM. In a subsequent pilot study involving five male mice undergoing similar surgical procedures, also with two sequential surgeries, but using xylazine as the alpha-2 adrenoreceptor agonist in the narcotic combination, none of the animals developed obstructive uropathy in the post-operative period or thereafter. We suggest that replacement of medetomidine with xylazine should be considered for general anesthesia in male mice undergoing multiple successive anesthesia events or in for procedures in older males.

## Introduction and results

The anesthetic combination of fentanyl, midazolam and medetomidine (FMM) is used widely for surgical procedures in rodents in lab settings. Is a preferred combination because it provides animals with thorough pain protection and stable general anesthesia during surgical procedures, it is well tolerated by rodents and, importantly, antagonists are available for all components meaning that the general anesthesia can be effectively and rapidly reversed.

However, in a recent cohort of 17 mice (5 male and 12 female, all C57Bl/6 mice modified to express cre in neurons in cortical layers 4 and 6^1^) undergoing multiple successive procedures under FMM general anesthesia over several weeks, 3 of the mice were found either deceased in their homecage or moribund and requiring immediate euthanasia on humanitarian grounds within the first 48 hours after recovery from a subsequent FMM anesthesia. In addition to these cases, we also report one additional case in which a mouse was found moribund after a single surgical procedure under FMM general anesthesia.

All of the adversely effected animals were male (none of the 12 female animals in the cohort had any observable adverse reactions after the anesthesia), and all were found to have dramatically distended bladders on postmortem necropsy. In two cases, more complete postmortem dissection of the urinogenital tract was performed, and both cases a solid blockage was found in the upper urethra. The animals in the main cohort all underwent two surgical procedures, a craniotomy and intracortical injection of AAV-GCaMP7f followed two to three weeks later by a second surgery to implant a light-weight implant for mounting a miniature microscope, opening of a craniotomy, including resection of the dura and installation of a cranial window for multiphoton imaging of neuronal activity. The full surgical procedures for both surgeries, including animal ethics declaration, have been published in Klioutchnikov et al. (2022)^1^. All these mice were without complication in the convalescence from the first surgery and showed no abnormal behavior or other complications in the period between the first and second procedures. Of the three animals suffering adverse reactions, one was found deceased and one moribund within 48 hours of the second surgical procedure, while the third was found moribund in its homecage within 48 hours of a subsequent general anesthesia for maintenance of the cranial window. Animal weight, duration of general anesthesia were all similar between the adversely effected and surviving males and between adversely effected males and females (Table 1, note that 1) statistics are not stated due to the limited sample size and 2) that the sex bias in the cohort resulted from deliberate selection of female mice for these experiments after detecting the vulnerability of male mice to the subsequent general anesthesia with FMM).

**Table 1.**
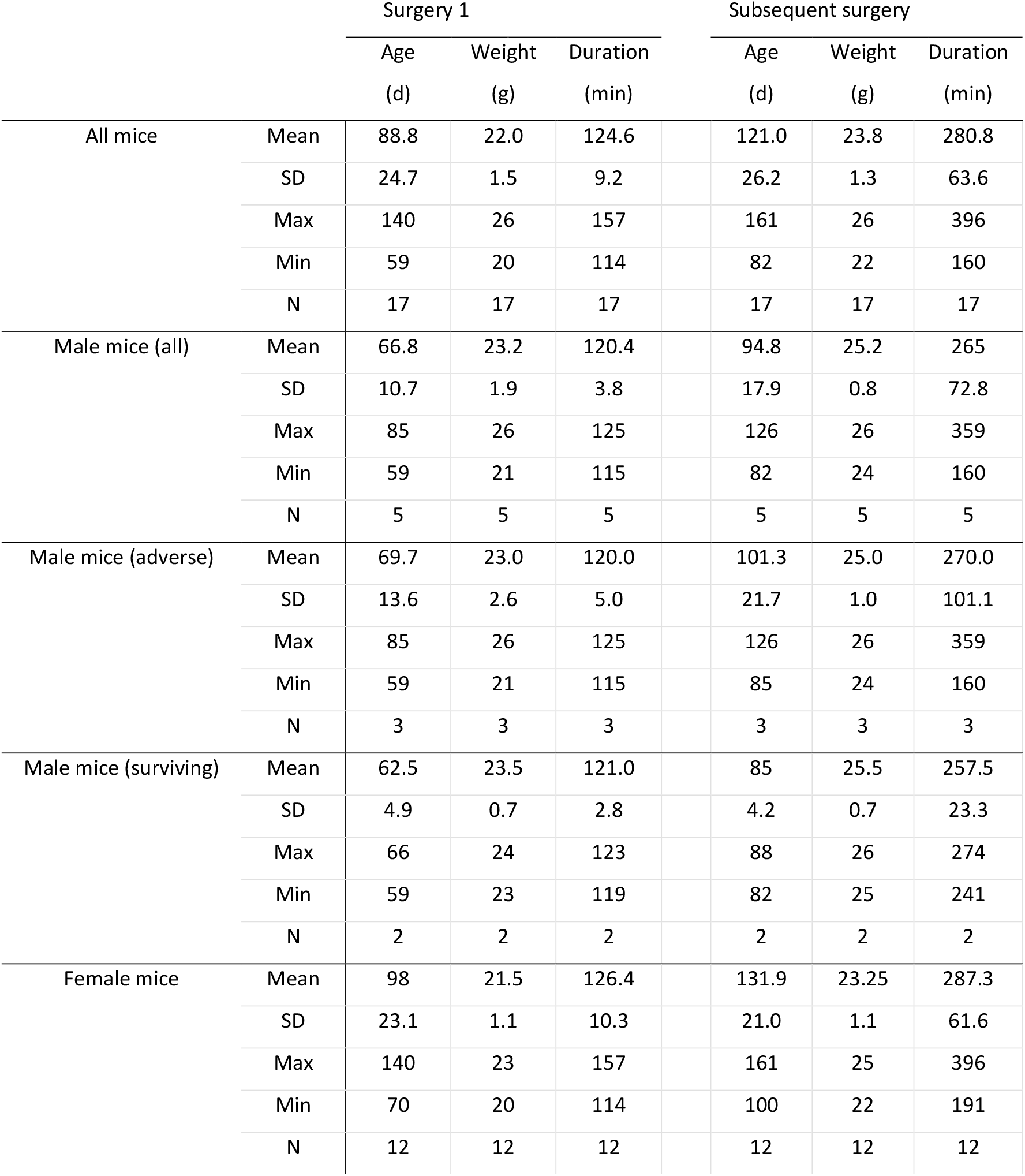
Animal characteristics for main animal cohort.

In addition to the cases described above, we also observed one additional case in which a male mouse was found moribund, also with a dramatically distended bladder, after a single anesthesia with FMM. In this case the animal had undergone only a single surgical procedure involving the opening of a small craniotomy for intracortical injection of cholera-toxin B, an anatomical tracer.

The ages of the adversely effected males were 85, 93 and 126 days prior to the surgery with the adverse event, while that of the surviving males were 82 and 88 days, and the animal with the adverse event after a single anesthesia was 210 days.

The presentation of the animals at the time of death was similar to that described by Wells et al. (2009)^2^, who reported similar observations in a group of male C57Bl/6 mice anesthetized with ketamine (50mg/kg) and medetomidine (0.5mg/kg). In this study, the authors performed a thorough postmortem work up on a subgroup of the effected animals, and concluded that this anesthetic combination is a potential risk factor for obstructive uropathy due to release of seminal coagulum. Urethral plugs, either naturally occurring, associated with infection or associated with specific strains or medical phenotypes have been previously reported to be associated with obstructive uropathy in male mice^3–10^, and another recent study has reported urethral obstructions associated with anesthesia using a drug combination including dexmedetomidine^11^, the active stereoisomer of medetomidine. Importantly, Wells et al. (2009) also reported that the morbidity and mortality they observed associated with anesthesia using ketamine and medetomidine was not observed after switching to an anesthetic combination using xylazine as a replacement for medetomidine.

We therefore conducted a pilot study using xylazine as an alternative alpha-2 adrenoreceptor agonist sedative in combination with fentanyl and midazolam for these surgeries to protect male mice from developing obstructive uropathy postoperatively. As for the experiments from which the cohort above were derived, the pilot study experiments were conducted in accordance with the institutional animal welfare guidelines of the Max Planck Society and with animal experimentation approval granted by the Landesamt für Natur, Umwelt und Verbraucherschutz Nordrhein-Westfalen, Germany (approval number 81-02.04.2020.A403). Experimental subjects were 5 male C57Bl/6 mice (4 *Ntsr1* (neurotensin receptor 1)-Cre mice and 1 wildtype). All animals were from the animal colony described in Klioutchnikov et al. (2022)^1^ from which the mice in the original cohort described above also came, with the *Ntsr1*-Cre mice showing only the Ntsr1-Cre phenotype and the wildtype no-Cre phenotype. For both the animals in this pilot study and the original cohort, the mice were housed in an SPF temperature- and humidity-controlled facility on a 12 h light–dark cycle with food and water available ad libitum. Mice were group-housed until surgery and singly housed afterwards. The animals were on average 189.4 ± 89.6 days old at the time of the first surgery (mean ± SD, range 112 to 328 days, N=5), with an average body weight of 31.0 ± 2.1 g (mean ± SD, range 29 to 34 min, N=5). The animals were anesthetized with the anesthetic combination fentanyl (50 μg/kg, Hameln pharma plus GmbH, Hameln, Germany), midazolam (5 mg/kg, Hameln pharma plus GmbH, Hameln, Germany) and xylazine (10 mg/kg, WDT, Garbsen, Germany) administered by i.p. injection, and then underwent the surgical procedure for intracortical virus injection, four being injected with AAV-GCaMP7f and one with the anatomical tracer cholera toxin B-subunit (Thermo-Fischer Scientific, MA, USA). All surgical, intra- and postoperative animal care and postoperative analgesia procedures were as described in Klioutchnikov et al. (2022)^1^. At the completion of the surgery, the animals were administered the antagonist combination naloxone (1.2 mg/kg, Ratiopharm, Ulm, Germany), flumazenil (0.5 mg/kg, Hikma, Amman Jordan) and atipamezole (5 mg/kg, Orion Pharma, Hamburg, Germany) via i.p. injection. After the induction dose, the animals reached surgical plane anesthesia after an average of 13.6 ± 10.1 min (mean ± SD, range 3 to 20 min, N=5). Average duration of the surgery was 114.4 ± 8.0 min (mean ± SD, range 104 to 124 min, N=5), with 3 animals requiring a single supplementary dose each of the anesthetic combination. After administration of the antagonist combination, animals recovered in an average of 2.4 ± 1.1 min (mean ± SD, range 1 to 4 min, N=5). After 1-7 weeks, the animals were again anesthetized with the same anesthetic combination described above, and underwent a second surgical procedure. Four underwent the surgical procedure for implantation of the microscope-mounting implant, craniotomy with durotomy and installation of a cranial window, while one had a second intracortical of an AAV encoding a cre-dependent red fluorescent protein (AAV1-hSyn-DIO-mCherry from Addgene, MA, USA) with the procedure being the same as that described above. As previously, all surgical, intra- and postoperative animal care and postoperative analgesia procedures as described in Klioutchnikov et al. (2022)^1^. At the time of this surgery the animals were an average of 217.8 ± 96.9 days old (mean ± SD, range 147 to 377 days, N=5), with an average body weight of 29.8 ± 2.2 g (mean ± SD, range 27 to 33 min, N=5). At the completion of the surgery, the animals were administered the antagonist combination described above. For this second surgical procedure, the animals reached surgical plane anesthesia after an average of 10.6 ± 3.4 min (mean ± SD, range 7 to 14 min, N=5) and average duration of the surgery was 231.8 ± 72.2 min (mean ± SD, range 112 to 303 min, N=5). During this time, one animal required a single supplementary dose of the anesthetic combination, two animals required three supplementary doses and two required four supplementary doses. After administration of the antagonist combination, animals recovered in an average of 5.0 ± 4.6 min (mean ± SD, range 1 to 10 min, N=5). Of these animals none suffered the adverse reaction, resulting in obstructive uropathy, described above, with all following an uneventful postoperative convalescence after both first and second surgical procedure.

## Discussion

Obstructive uropathy is a condition that has been widely reported to effect male mice^3–10^, and which can, in chronic cases, cause distress, discomfort, reduced breeding capacity and self-inflicted damage to the genital region, and in acute cases can cause complete block of the urethra with fatal consequences. Multiple reports, including the current case study, now document an association between anesthesia with anesthetic combinations involving the alpha-2 adrenoreceptor agonist medetomidine and the development of obstructive uropathy in the days after recovery from the anesthesia^2, 1^. In previous experimental cohorts where the FMM combination was used for general anesthesia in male mice we did not observe any cases of obstructive neuropathy on recovery from anesthesia^12^. The most pertinent difference between the previous cohort and the current one is that in the previous cohort the animals underwent only a single surgical procedure, and were younger at the time that they were administered the dose of FMM. In all but one case where we observed adverse reaction to the FMM combination the animals were recovering from a second general anesthesia at the time of the adverse reaction, and were older than 85 days at the time of the surgery. The one animal suffering an adverse reaction after a single general anesthesia with FMM was 201 days at the time of the procedure, suggesting that older animals are more susceptible to obstructive uropathy, though the sample size is clearly too small to make a conclusive statement. We have also never observed obstructive uropathy in male rats^13^, even though they were involved in very similar experimental cohorts to those described here where the adverse reactions occurred in male mice. As demonstrated previously, we also found here that obstructive uropathy was not observed after replacement of medetomidine with xylazine (10 mg/kg). We suggest that the combination of fentanyl, midazolam and xylazine is an effective, stable and well tolerated combination for general anesthesia in male mice, and that it should be considered both for procedures on older male mice as well as in cases involving multiple procedures under general anesthesia.

## Methods

The experiments were conducted in accordance with the institutional animal welfare guidelines of the Max Planck Society and with animal experimentation approval granted by the Landesamt für Natur, Umwelt und Verbraucherschutz Nordrhein-Westfalen, Germany (approval number 81-02.04.2020.A403).

### Anesthesia and antagonist combinations for xylazine test group

Experimental subjects were 5 male C57Bl/6 mice (4 *Ntsr1* (neurotensin receptor 1)-Cre mice and 1 wildtype). The mice were anesthetized with the anesthetic combination fentanyl (50 μg/kg, Hameln pharma plus GmbH, Hameln, Germany), midazolam (5 mg/kg, Hameln pharma plus GmbH, Hameln, Germany) and xylazine (10 mg/kg, WDT, Garbsen, Germany) administered by i.p. injection. At the completion of the surgery, the animals were administered the antagonist combination naloxone (1.2 mg/kg, Ratiopharm, Ulm, Germany), flumazenil (0.5 mg/kg, Hikma, Amman Jordan) and atipamezole (5 mg/kg, Orion Pharma, Hamburg, Germany) via i.p. injection.

### Anesthesia and antagonist combinations with original anesthetic combination

Experimental subjects were 17 mice (5 male and 12 female), all C57Bl/6 mice modified to express cre in neurons in cortical layers 4 and 6. Animals were anaesthetized with a three-component anesthetic cocktail (3K) consisting of fentanyl (50 μg/kg, Hameln pharma plus GmbH, Hameln, Germany), midazolam (5 mg/kg, Hameln pharma plus GmbH, Hameln, Germany) and medetomidine (0.5 mg/kg, Zoetis, NJ, USA). At the completion of the surgeries the animals were administered an antagonist combination consisting of naloxone (11.2 mg/kg, Ratiopharm, Ulm, Germany), flumazenil (0.5 mg/kg, Hikma, Amman Jordan) and atipamezole (0.75 mg/kg, Orion Pharma, Hamburg, Germany).

### Animals

Mice were housed in an SPF temperature- (21 ± 1°) and humidity-controlled (>45%) facility on a 12 h light–dark cycle with food and water available ad libitum. Mice were group-housed until surgery and singly housed afterwards.

The mice *(Mus musculus*) used in this study resulted from crossing heterozygous animals from two Cre-expressing lines, Ntsr1-Cre mice and Scnn1a-Cre mice. *Ntsr1* (neurotensin receptor 1)-Cre mice (B6.FVB(Cg)-Tg(Ntsr1-cre)Gn220Gsat/Mmcd) were obtained from the Mutant Mouse Resource and Research Center (MMRRC, #030648-UCD) and donated by Nathaniel Heintz (The Rockefeller University, GENSAT, New York, NY, USA) and Charles Gerfen (National Institutes of Health, National Institute of Mental Health, Bethesda, MD, USA), with Cre recombinase predominantly expressed in layer 6 of the cortex^14^. *Scnn1a* (sodium channel, nonvoltage-gated 1 alpha)*-Cre* mice ([Tg(Scnn1a-cre)3AibsTg(Scnn1a-cre)3Aibs](http://www.informatics.jax.org/accession/MGI:3850183)) were obtained from the Jackson Laboratory (#009613) and donated by Ed Lein (Allen Institute for Brain Science, Seattle, WA, USA) and Theresa Zwingman (Allen Institute for Brain Science, Seattle, WA, USA) with Cre expression in cortex layer 4^15^. The Cre-animals were maintained in a heterozygous state with C57BL/6J background. Expression patterns of these transgenic Cre lines have been described in the original papers cited.

### Surgical procedures for fluorescent labelling of neurons with jGCaMP7f

Prior to surgery, all instruments were sterilized by autoclaving, and the glass injection capillaries were sterilized by heat sterilization. Animals in the xylazine test group were anesthetized as described in the section “*Anesthesia and antagonist combinations for xylazine test group*” above, and the animals in the original cohort were anesthetized as described in the section “*Anesthesia and antagonist combinations with original anesthetic combination*” above. Body temperature was maintained at 37–37.5 °C with a heating pad and heater controller (FHC, ME, USA). Animal status and depth of anesthesia was monitored approx. every 15–30 min, and anesthesia maintained throughout with supplementary doses of 30–80% of the anesthetic combination given as necessary in order to maintain absence of withdrawal and corneal reflexes. The animals were then placed in a stereotaxic apparatus, the hair on the scalp removed and the skin cleaned with 70% ethanol. The right parietal bone was exposed and a burr hole drilled (approx. 500 μm diam.) at 3.0 mm posterior and 2.6 mm lateral relative to bregma. A small slit was made in the dura underlying the burr hole, and a glass injection capillary with a beveled tip containing a high-titer solution of AAV1/2 coding for Cre-dependent jGCaMP7f (AAV1/2.hSyn.FLEX.jGCAMP7f, Addgene, MA, USA) was advanced posteriorly into the cortex approx. 1650 μm at an angle of 25° relative to horizontal along a trajectory parallel with the sagittal suture. An injection of approx. 180 nL of the virus solution was made with a nanoliter injection device (Nanoject II, Drummond Scientific Company, PA, USA). After a delay of 5 min, the tip of the capillary was withdrawn approx. 240 μm, and an injection of approx. 115 nL made. After another delay of 5 min, the tip of the capillary was again retracted approx. 240 μm, and another injection of approx. 115nL made. After a final waiting period of 5 min, the glass capillary was slowly withdrawn, the craniotomy covered with silicone (KwikSil, WPI, FL, USA) and the skin sutured closed using 5/0 vicryl sutures (Ethicon, NJ, USA). Animals were then administered buprenorphine (30 μg/kg, Bayer, Leverkusen, Germany) and carprofen (5 mg/kg, Zoetis, NJ, USA) for post-operative analgesia, and then the respective antagonist combinations described in the sections “*Anesthesia and antagonist combinations for xylazine test group*” and “*Anesthesia and antagonist combinations with original anesthetic combination*” above.

### Second surgical procedure for installation of a cranial window for imaging

All surgical instruments and solutions used were autoclaved prior to commencement of the procedures described below. Three to five weeks after the surgery to label neurons with jGCaMP7f, animals in the xylazine test group were anesthetized as described in the section “*Anesthesia and antagonist combinations for xylazine test group*” above, and the animals in the original cohort were anesthetized as described in the section “*Anesthesia and antagonist combinations with original anesthetic combination*” above. Body temperature maintained at 37–37.5°C. Animal status and depth of anesthesia monitoring procedures were as described in the section on labelling neurons with jGCaMP7f described above. Anesthesia was maintained with supplementary doses of 30–80% of the anesthetic combination. The hair on the dorsal aspect of the skull was removed and the skin cleaned with 70% ethanol. A midline incision in the skin over the parietal bones was made, the skin retracted and galea removed to expose the parietal bones, including the site of the previous burr hole. The exposed bone was then cleaned with hydrogen peroxide solution (3% by volume in sterile saline) and thoroughly washed with sterile saline. The bone was then mechanically roughened prior to application of a layer of dental adhesive (Optibond, Kerr, CA, USA). A custom-made headplate (described in full in^1^) was then fixed to the skull over the Optibond layer with dental composite (Charisma, Kulzer GmbH, Hanau, Germany). The central aperture was placed such that the burr hole from the previous surgery was located approximately centrally in the medial-lateral axis and near the anterior edge of the aperture. The skin incision was then closed firmly around the headplate using 5/0 vicryl sutures (Ethicon, NJ, USA). A circular craniotomy with a diameter of approx. 3.5 mm was then opened in the center of the headplate aperture, including at the anterior margin of the site of the previous craniotomy. The dura was then removed and the cranial window closed using a pre-formed plug and coverslip (circular, 5 mm diam., 100 μm thickness, CS-5R-0, Warner Instruments Holliston, MA, USA; plug custom pre-formed as a 300 or 400 μm tall cylinder of KwikSil silicone in the center of the circular coverslip after the description in^16^). Animals were then administered buprenorphine (30 μg/kg, Bayer, Leverkusen, Germany) and carprofen (5 mg/kg, Zoetis, NJ, USA) for post-operative analgesia, and then the respective antagonist combinations described in the sections “*Anesthesia and antagonist combinations for xylazine test group*” and “*Anesthesia and antagonist combinations with original anesthetic combination*” above.

## Acknowledgements

We would like to thank Nadine Schulz for useful discussions. Funding from the Max Planck Society.

## Data Availability

Data available upon request.

